# Genetic insights into Iron Age Saka culture: Ancient DNA analysis of the Boz-Barmak burial ground, Kyrgyzstan

**DOI:** 10.1101/2025.11.03.686206

**Authors:** Aigerim Rymbekova, Pere Gelabert, Alejandro Llanos-Lizcano, Kaaviya Balakrishnan, Michelle Hämmerle, Olivia Cheronet, Aida Abdykanova, Keldibek Kasymkulov, Michelle Hrivnyak, Jacqueline T. Eng, Ron Pinhasi, Martin Kuhlwilm

## Abstract

The nomadic cultures of the Iron Age played an important role in shaping the genetic and cultural landscape of Eurasian populations. Yet despite its key geographical location, the Central Eurasian region remains underrepresented in ancient DNA studies of humans. We address this gap through genomic analysis of 12 individuals from the Boz-Barmak burial site in Kyrgyzstan associated with Saka pastoralists (4th-2nd centuries BCE), 9 of which yielded low-coverage genomes. Genetic clustering analysis placed these individuals within the genetic variation of ancient and modern Central Eurasian and Siberian populations. We found no evidence of first-degree relatives in a kinship analysis, however a network of second- and third-degree relationships seems to be present. Notably, all male individuals share the same Y-chromosomal haplotype, common in present-day Kyrgyz and Tajik groups, while mitochondrial DNA showed comparably high diversity, with distinct haplogroups observed across the analysed individuals. These findings are in line with archaeological and ethnographic evidence of patrilocality in Early Iron Age Saka, where male lineages remained stable across generations, while female mobility contributed to genetic diversity. Our study complements our understanding of the interplay between kinship, social organization and population history in nomadic cultures.

## Introduction

In the Eurasian Steppe, a dynamic population history with continental-scale migration events took place, with Central Eurasia playing a crucial role as a crossroads of continents, as demonstrated by the westward expansion of pastoralist groups from the Altai region around 5,000 years ago. These migrations into Europe and South Asia are believed to have spread some of the Indo-European languages, fundamentally reshaping the linguistic and genetic landscape of a large part of the world [1–5].

The Iron Age (IA) (ca. 1000 BCE – 500 CE) marks an important period in Eurasian history, when nomadic horse-riding cultures emerged as powerful forces that reshaped the genetic and cultural landscape of populations across the continent. Among these groups, the Saka peoples, closely associated with the Scythians, stood out as one of the most prominent IA cultures. Their presence across the vast Central Eurasian Steppe, inhabiting various ecological and climatic conditions, led to the development of distinct economic systems. This resulted in the profound complexity of the Saka confederation [6, 7].

It is believed that Saka were a political union of different tribes united by military discipline and at the core of their society stood the kin, an extended unit composed of several families [8]. These kin groups could hold territories, with land regarded as communal property controlled by kin leaders. Some kins were noble houses. Among nomads, the principle of kin-based inheritance of supreme power was rooted in the idea that authority belonged to the entire ruling lineage [9]. According to the genetic evidence, the Saka people likely originated in the Central Eurasian Steppe and were descendants of Late Bronze Age (BA) steppe herders admixed with hunter-gatherers from neighboring regions [10-13]. The differences in the sources and proportions of hunter-gatherer ancestry among the people associated to the Saka culture show that groups of the confederation arose from separate local populations in different places [14, 15]. Notably, in some populations, such as Saka of present-day Kyrgyzstan, Iranian Neolithic ancestry is also present [4], suggesting interaction with the Bactria-Margiana Archaeological Complex (BMAC), a sophisticated sedentary civilization of farmers in present-day Turkmenistan [16]. While these studies provided insights into the origins of Saka, their genetic substructure and kinship practices remain poorly understood.

The iconic and often massive monuments of the steppe, kurgans, were widespread from the northern Black Sea in the west to the region of Siberia and northern China in the east, including Central Eurasia. Archaeological evidence suggests that the oldest dated Scythian kurgans are from the 9^th^ century BC found in present-day Tyva republic, southern Siberia [17]. The size of the Scythian kurgans and graved artifacts, including various weapons, jewelry and pottery, appear to be correlated with the social status and degree of nobility of the person buried [6]. For example, at the Besshatyr kurgan complex in Kazakhstan, the mounds of Saka military leaders reached up to 105 meters in diameter and 17 meters in height. In Scythian-related cultures, including Saka, the burials were arranged in straight-line sequences, or meridional chains, that varied in size and complexity, reflecting the considerable, large-scale communal effort invested in their construction and the existence of social hierarchy [18]. The burials of armed warrior Saka women suggest that, despite the prevailing patriarchal system, women could hold important positions within the community, could have been free to choose their husbands and, equally with men, took part in military activities [19, 20]. Nevertheless, the fine-scale social structure of the Saka still remains a subject of debate.

To address questions on genetic diversity and social structure of the Sakas, here we focus on the Boz-Barmak burial ground in present-day Kyrgyzstan. Our approach combines archaeological analysis of burial structures and artifacts with ancient DNA analysis, to enhance our understanding of Saka culture.

## Results

### Burial ground archaeological context

In 2022, as a result of rescue excavations, the Boz-Barmak burial ground was excavated during the construction of the Balykchi–Ak-Olon railroad. The Boz-Barmak burials are located in the Boz-Barmak mountains, approximately 4 km southeast of Balykchi town, along the southwestern shore of Issyk-Kul Lake. The burial ground can be dated over a period between the 4th-2nd centuries BC, which chronologically fits within the Iron Age.

The site consists of 14 single-individual burials aligned in a chain from north to south oriented by head to western sector, though some are positioned outside the main sequence (Fig.1). The burial mounds have diameters from 7 up to 10–14 meters and reach a height of about 1 meter, indicating that they belonged to the local elite. The burials exhibit a sophisticated construction. The burial pits were covered with timber placed transversely, followed by a layer of shrubs. This structure was then secured and surrounded by clay blocks, which were cut from nearby sources to form the base of the mound. The clay base was reinforced with layers of rock. Additionally, stone rings were placed around the perimeter of the burials to stabilize the mound.

**Figure 1.**
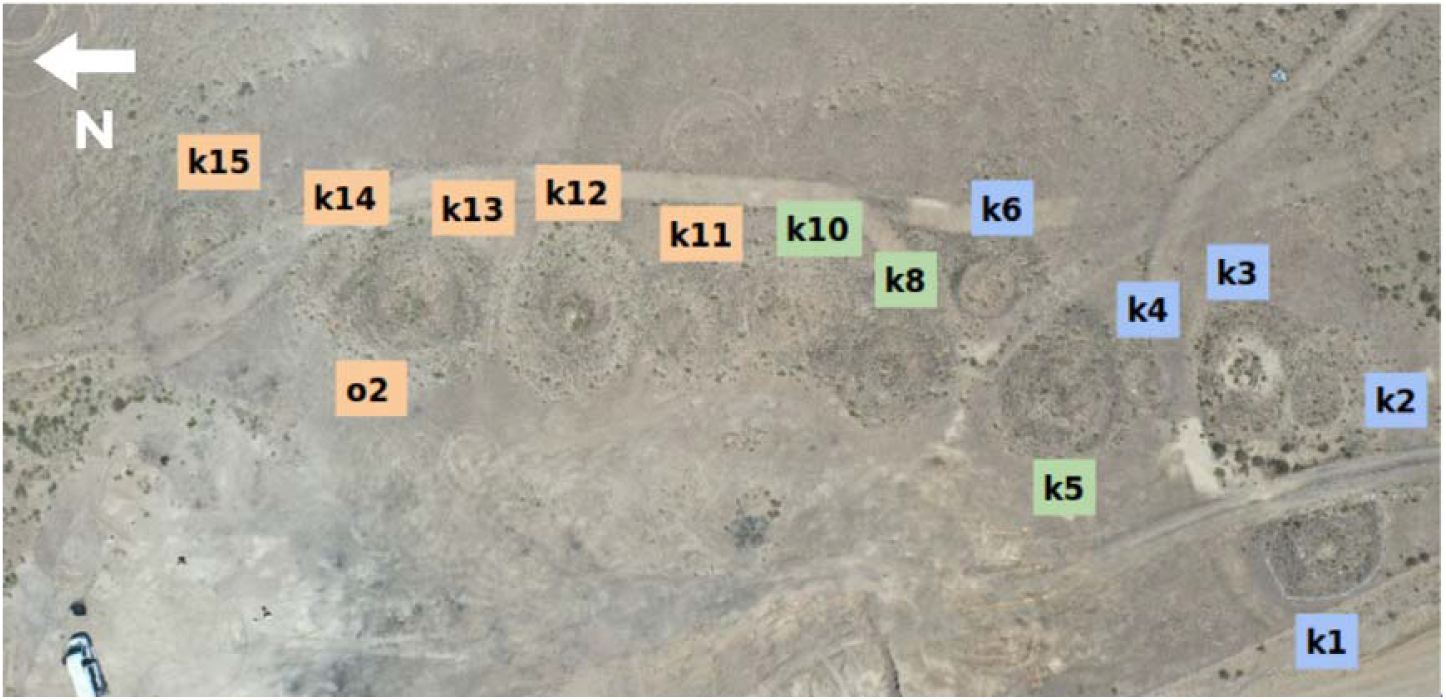
The site consists of 14 burials aligned in a chain from north (indicated with a white arrow) to south. The map shows a possible chronological sequence of mound construction, based on the current interpretation, with kurgans grouped into three clusters from earliest to latest: green cluster (K10, K8, K5), orange (K11, K12, K13, K14, K15, Object 2) and blue (K6, K4, K3, K2, K1).

Analysis of the material culture found in the burials indicates that the burial chain was not organized in a strict north-south orientation but followed a more irregular pattern, probably from center to periphery. Furthermore, our analysis of cultural artifacts and mound construction suggests that the kurgans were built sequentially, from earliest to latest, and may be grouped into three clusters (the following are kurgan numbers): 1) 10.1, 10.2, 8, 5; 2) 11, 12, 13, 14, 15 and object 2; 3) 6, 4, 2, 1 and 3. Kurgan 10 (K10) is proposed as the earliest burial, based on the presence of a rhombus-shaped golden plate decorated with a trefoil motif, as well as a reconstructed handled jar, characteristic of the pre-Hellenistic period.

Kurgans 5, 8, 10, 11, 12, and 13 are larger in size and appear to be part of the main chain. Kurgans 1, 4, 5, 6, 8, 15 have northern central niches to place pottery set with ritual food. The individual in Kurgan 13 (K13) is interpreted as female, based on the discovery of a gold ring earring with a small soldered ring, similar to findings in kurgans 11 and 10.2, all are females. Later analysis did not yield enough genetic material of the individual in K13 to corroborate archaeological evidence of sex. Kurgans 6, 14 and 15 are smaller in size and appear not to represent leaders of the kin, but rather individuals who played crucial supporting roles in the leadership structure. Kurgan 6 is located outside the main burial chain and has a smaller size. However, it features an outstanding burial mound structure and has been found in situ. In total, only four burials were discovered in situ — Kurgans 4, 6, 13 and object 2. The other burials are presumed to have been looted. Kurgan 6 has two rings supporting the mound and a soft filling in the center. The inner ring was originally a separate mound, composed of rock-covered clay blocks. It closely resembles K13, as both feature two vertically standing rocks in the outer ring, oriented in an east-west direction. The space between these two rocks is enclosed by another rock positioned horizontally. The burial contains unusual artifacts, including an ancient derivative bone buckle adorned with plant motifs, a later-period braid fixator or part of headdress decorated with gold plates, and a small bronze folding mirror with an umbo. Kurgans 9, 4, 2 and object 2 are satellite burials positioned near kurgans 8, 13, 5 and 3: specifically in the northwestern corner with object 2 and K9, and in the southern side with kurgan 4 and 2. Kurgan 9 consists of two burials, arranged one beneath the other. Kurgan 4 is small in size and attached to K5 in the south. The individual buried in K4 was young and maybe considered as son and heir of the individual in K5. Kurgan 2 is attached to K3.

Object 2 is a burial without a mound, consisting of a grave pit with shoulders covered by timber and elongated rocks. The buried woman wore a wooden headdress with rigid outlines, resembling a helmet with holes in the upper and occipital parts. A bronze pin was found vertically positioned in the upper hole of the headdress, and a single bronze earring was discovered next to her left ear. Additionally, the discovery of a horse coccyx bone in the burial, likely as part of ritual food suggests that she held a high social status, signifying respect for her age and position in society.

Among the burials, only object 2 and kurgans 11, 12, and 13 had grave pits with shoulders. These pits were unusually deep, over 2 meters in the case of kurgans 11, 12 and 13, allowing enough space to lower the deceased inside. The original pit was larger, and at a depth of about 1 meter, another pit was dug in the center, leaving shoulders or ledges around its edges. All the deceased were buried in wooden-frame coffins, possibly covered with felt. Burial pits of K10.2 and K11 contained more elongated coffins, extending up to one meter beyond the head. This extra space may have been used for storing artifacts, which were later probably looted. Despite the looting, the upper part of the woman buried in K11 remained in situ. The preserved shape of a headdress was found on her head, and beneath it, a bronze buckle in the form of a mirror and a bronze pin were discovered. Only women in object 2 and K11 have preserved headdresses. In the mound of K12, scattered remains of the upper body and head were discovered. The primary burial of K12 was found in situ without any material objects.

### Ancient DNA analysis

We extracted DNA from 12 individuals, following standard ancient DNA procedures. Sequencing libraries were shotgun sequenced, and mitochondrial DNA (mtDNA) capture was performed (Methods). Two individuals were excluded from deeper sequencing due to low coverage after mitochondrial capture (BB.K.4, BB.K.8), indicating poor DNA preservation (Table S1). We aimed to obtain approximately 1-fold genome-wide coverage through sequencing, resulting in coverage between 0.1-fold and 1.4-fold (Table 1). One individual (BB. K.10.2) with low coverage (0.03-fold) and substantial contamination (16.7%) was removed from further analyses, leaving 9 individuals. Contamination estimates for other samples range between 1.7% and 5.1%, which falls within the expected range for ancient DNA samples. We find typical patterns of ancient DNA damage (Fig S1), and endogenous DNA estimates range from 3% to 67.2%. We estimated the coverage on the autosomes and the sex chromosomes separately, which enables us to assign the sex through their respective ratios. Among the nine individuals included in further analyses, six were determined as females and three as males (Table 1).

**Table 1.** Basic metrics on the quality of the samples and genetic sex determination. (Endog. = endogenous DNA; Modern cont. = modern contamination; Auto cov. = autosomal coverage; X cov. - X chromosome coverage; Y cov. = Y chromosome coverage)

| ID | Total reads | Endog.% | Modern cont.% | Auto. cov. | X cov. | Y cov. | Sex |
| --- | --- | --- | --- | --- | --- | --- | --- |
| BB.K.1 | 91,576,637 | 67,2 | 2,5 | 1,4 | 1,43 | 0,01 | F |
| BB.Obj.2 | 105,353,653 | 24,7 | 2,1 | 0,6 | 0,62 | 0,01 | F |
| BB.K.6 | 122,390,978 | 20 | 1,7 | 0,6 | 0,31 | 0,08 | M |
| BB.K.9 | 169,442,108 | 3 | 2,0 | 0,1 | 0,05 | 0,01 | F |
| BB.K.10.1 | 102,344,280 | 23,2 | 5,1 | 0,5 | 0,28 | 0,07 | M |
| BB.K.10.2 | 92,365,334 | 1,5 | 16,7 | 0,03 | 0,03 | 0 | F |
| BB.K.11 | 85,216,391 | 66,3 | 1,7 | 1,4 | 1,45 | 0,01 | F |
| BB.K.12 | 120,774,880 | 32,9 | 5,1 | 0,6 | 0,62 | 0,01 | F |
| BB.K.14 | 140,856,045 | 13,2 | 3,0 | 0,7 | 0,35 | 0,09 | M |
| BB.K.15 | 119,966,700 | 27,7 | 2,1 | 0,8 | 0,62 | 0,01 | F |

We explored the biological relationships within our dataset with ancIBD [21], finding no first-degree relatives. However, we found three pairs of individuals with a second degree of relatedness and three pairs with third degree (Fig. 2A), according to shared IBD fragments longer than 12 centimorgans (cM). We propose the second degree relationship to be half-siblings, grandparent/grandchild or aunt/uncle/ niece/nephew on the paternal side. Only negligible shared IBD fragments were found on the X chromosome. We also explored inbreeding within our dataset using hapROH [22] by analyzing co-inherited identical haplotypes, or runs of homozygosity (ROH). Most individuals had no detectable ROH, likely indicating absence of close kin unions. Based on the amount of ROH and the number of segments found in three other individuals, we suggest that the parents of those individuals could have been fourth or fifth cousins (Table S2, Fig. S2). We do not find closely related individuals in previously published BA and IA Central Eurasian populations (Fig. S3).

**Figure 2.**
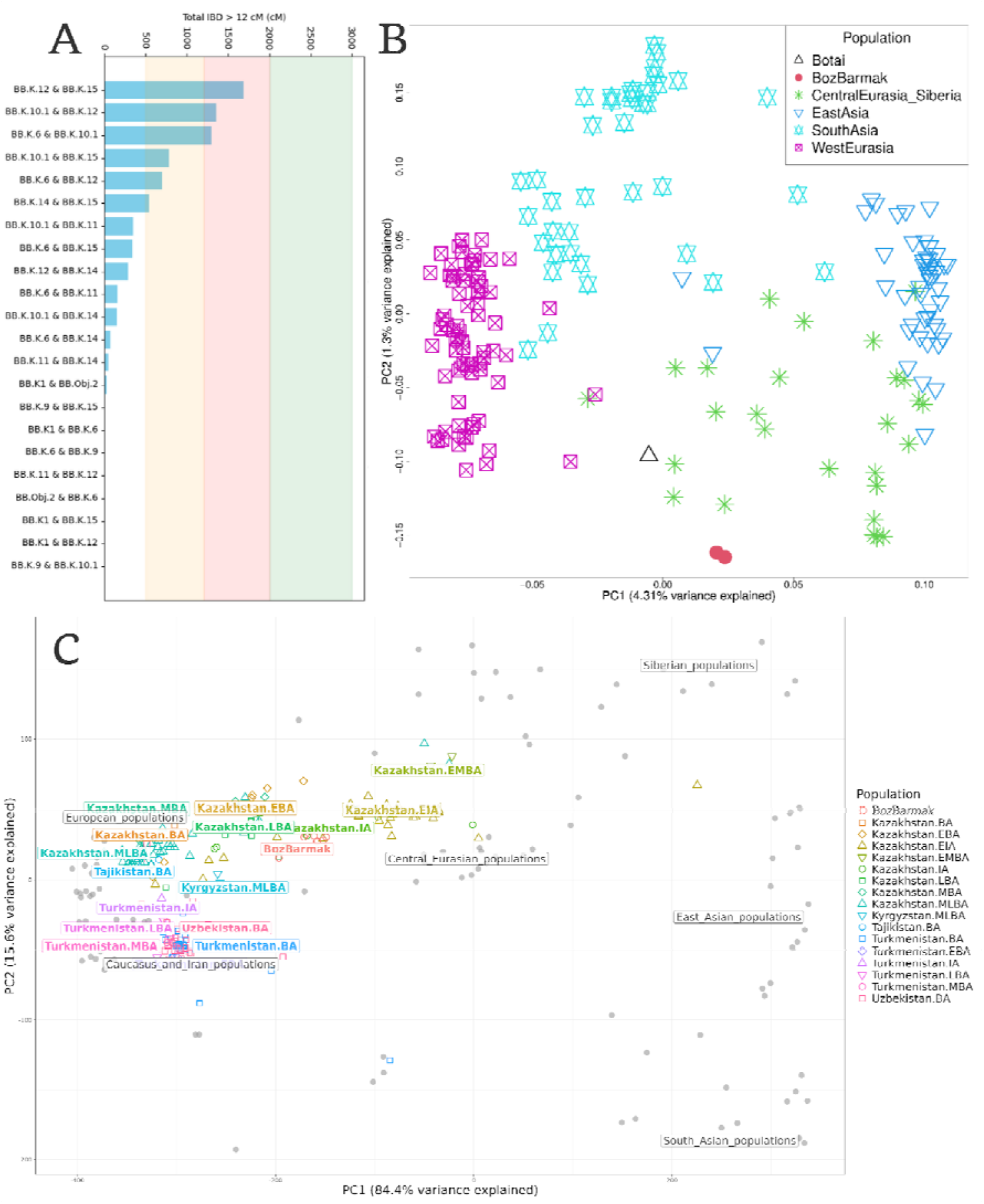
A. Pairwise Identity by Descent (IBD) sharing in centimorgans (cM) with relationship threshold ranges: first degree range colored green, second degree - pink, third degree - yellow, higher degree – white. B. Principal Component Analysis of Eurasian diversity with two individuals from Boz- Barmak, BB.K.1 and BB.K.11, based on whole-genome sequencing data. The reference populations include modern populations from Western Eurasia, Central Eurasia, Siberia, South Asia and East Asia, as well as a Bronze Age Kazakh Botai individual. C. Principal Component Analysis of Bronze Age and Iron Age Central Eurasian populations and Boz-Barmak individuals, based on genome-wide captured data for common human variation, in relation to the modern Eurasian reference panel shown in gray. (BA = Bronze Age, EBA = Early Bronze Age, MBA = Middle Bronze Age, MLBA = Middle Late Bronze Age, LBA = Late Bronze Age, IA = Iron Age, EIA = Early Iron Age)

We performed a Principal Component Analysis (PCA) to assess the genetic relationship between Boz-Barmak individuals and broader Eurasian populations. Using a reference panel of whole genome data of ancient and modern Central Eurasians, we analyzed BB.K.1 and BB.K.11, the two individuals with the highest coverage. These individuals cluster closely together and generally close to the variation of Central Asia and Siberia (Fig. 2B). We then performed a PCA analysis using only BA and IA populations from Central Eurasia, based on data obtained using targeted capture, intersected with genotype calls at the target variants across all nine Boz-Barmak individuals. The PCs were computed using modern Eurasian individuals, while the Boz-Barmak individuals together with BA and IA populations were projected onto the top two PCs. The Boz-Barmak individuals exhibited genetic affinity in particular to IA individuals from Kazakhstan (Fig. 2C). These results were corroborated with outgroup f3 statistics (Fig. 3A): the test f3(Boz-Barmak; Ancient Eurasian population, Mbuti) showed the strongest shared genetic drift between Boz-Barmak individuals and Kazakh populations.

**Figure 3.**
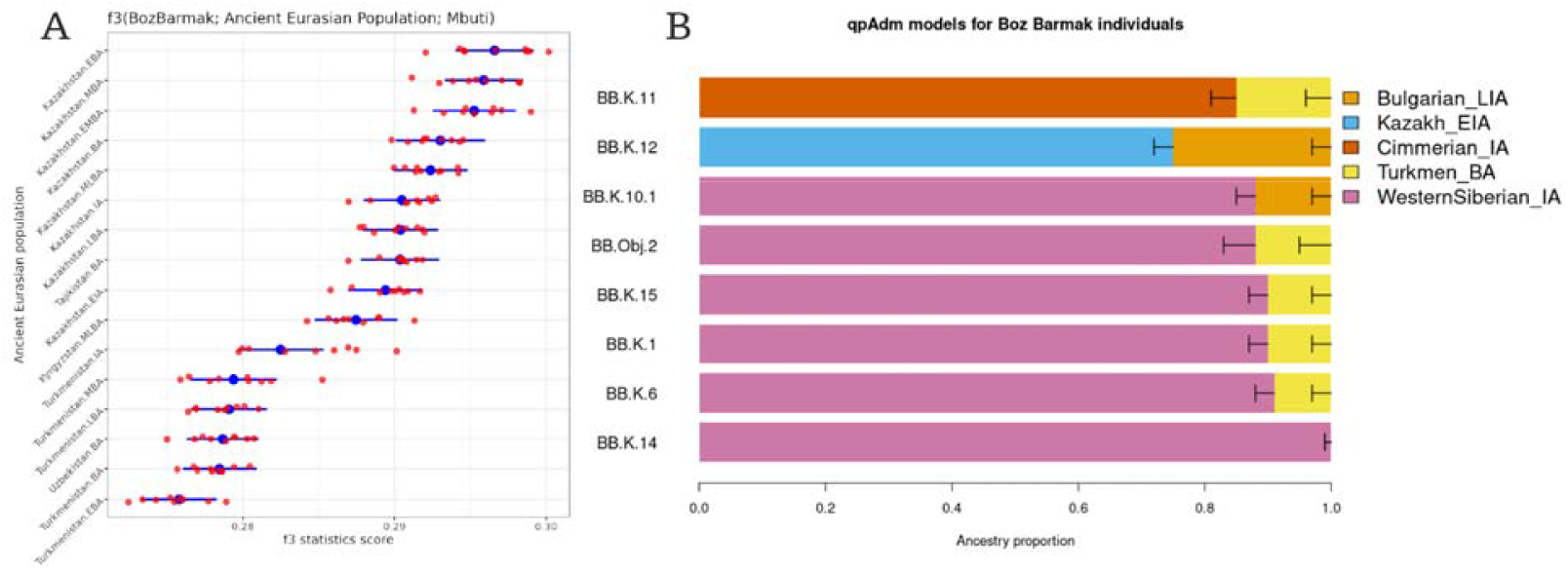
A. The f3-statistics score distribution of Boz-Barmak individuals with Bronze Age and Iron Age Central Eurasian populations. The blue dots represent per population values with standard error intervals and the red dots are per individual values. (BA = Bronze Age, EBA = Early Bronze Age, MBA = Middle Bronze Age, MLBA = Middle Late Bronze Age, LBA = Late Bronze Age, IA = Iron Age, EIA = Early Iron Age). B. Ancestry proportions based on qpAdm analysis. The proportion of each source population is represented by the box on the x axis. Horizontal bars represent ±1 standard error estimated by qpAdm using 5-cM block jackknifing.

To investigate potential ancestral sources and admixture proportions, we modeled our individuals using qpAdm [23]. We tested one-way models with a single source using BA and IA populations of the Pontic-Kazakh Steppe. One individual (BB.K.9) with the lowest genome-wide coverage (Table 1) was excluded from the analysis due to a low number of sites overlapping with the reference populations, as the threshold was set to 100,000 sites. Our analysis revealed that 6 out of 7 individuals included in the analysis can be successfully modeled using a one-way model with an Eastern Scythian-related IA sedentary culture – Sargat from Western Siberia (WestSiberian_IA). Previous genomic studies showed that the Sargat people descended from a gene pool that formed as a result of mixture between western Scythian groups and a local Eastern Eurasian population with a smaller amount of gene flow from an Iranian-related source [3, 16, 24]. Based on the archaeological evidence, Sargat culture is believed to be a geographical contact zone between those peoples [25, 26]. Similarly, 3 individuals out of 7 can be modeled using a one-way model with pre-Scythian IA nomads from Pontic-Caspian Steppe – Cimmerians (Cimmerian_IA). Genomic analyses have demonstrated that they had originated in the Far East and had an east-west admixture gradient across the Eurasian Steppe with significant amounts of ancestry from Near Eastern populations [15, 27-29]. One individual’s genetic profile (BB.K.11) could not be adequately modeled by the one-way models. To investigate further, we used two-way models to analyze Boz-Barmak individuals as a genetic mixture between two potential ancestry sources. The qpAdm results indicate that all 7 individuals can be modeled having either Cimmerian, Sargat or a pre-Saka IA nomad culture from central and north Kazakhstan, Tasmola (Kazakh_EIA), as a primary genetic ancestry component. Notably, a two-way admixture model involving 84% Cimmerian_IA and 16% Turkmenistan_BA yielded a good fit for the BB.K.11 individual (p-value=0.358, stderr=0.04). Other individuals were successfully modeled with several two-way models, including WestSiberian_IA (85-95%) and Turkmenistan_BA/Bulgaria_LIA (5-15%) or Kazakh_EIA (75-80%) and Tajikistan_BA/Bulgaria_LIA (20-25%). The best working models are shown in Figure 3B, and detailed results are provided in Supplementary Table S3. These results demonstrate heterogeneity within the Saka and, in line with previous findings, show that the gene pool of IA Saka in present-day Kyrgyzstan is a mixture of different western and eastern Eurasian sources. Furthermore, we find an affinity to southern populations related to Neolithic Iranians and the Mesolithic Caucasus hunter-gatherers.

The mtDNA analysis resulted in assigning each individual a different haplogroup, all of which are in line with broadly Eurasian ancestry (e.g. Yakut C4a, Uzbek HV6, Chinese K2a, Eastern Europe U2e). The male individuals were all assigned to one Y chromosome haplogroup, R1a1a1b2, which is commonly found in present-day Kyrgyz and Tajik individuals (Table 2).

**Table 2.**
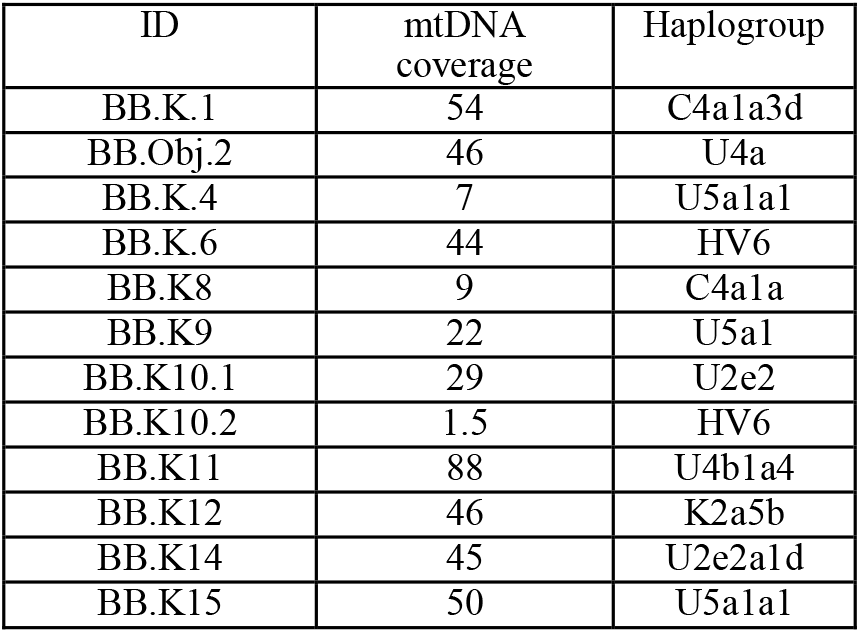
Mitochondrial DNA coverage and haplogroup assignment.

Finally, we screened the sequencing data for genetic traces of potential pathogens. Although we observed matching reads for multiple potentially pathogenic lineages, we did not obtain strong evidence for specific viral, bacterial or fungal infections in these individuals (Table S4).

## Discussion

In this study, we presented genetic data from 12 Early Iron Age pastoralists from the Boz-Barmak burial site in present-day Kyrgyzstan, with 9 low-coverage whole genomes (on average 0.7-fold coverage). The Boz-Barmak individuals were shown to be of Central Eurasian ancestry and are best modeled as a mixture between EIA Scythian-related groups, such as the Cimmerians and Sargat culture, and an Iranian-related source. This component includes ancestry from Neolithic pastoralists from South Central Asia, Turkmenistan and Tajikistan, and Mesolithic Western Eurasian hunter-gatherers from Bulgaria. These results corroborate historical hypotheses suggesting a cultural connection between the northern steppe populations and southern civilizations. Our findings support the genetic heterogeneity of the Saka by demonstrating several possible primary ancestry source populations and various admixture proportions among the Boz-Barmak individuals.

By integrating ancient DNA analysis results with archaeological data, we gained insights into the social structure and kinship practices of Early Iron Age Saka people at Boz-Barmak. The genetic evidence revealed that the burial ground was organized around a network of second-, third- and higher-degree genetic relationships. Crucially, the presence of diverse ancestral backgrounds within this network suggests that the social concept of kinship could have been flexible enough to tie individuals from varied lineages into a single kin group or extended family. The group might have been local elite or individuals closely associated with them as the burial chain consists of large and complex kurgans with valuable material artifacts. Archaeological evidence indicates that such chains were carefully planned [30, 31], a pattern also evident at Boz-Barmak, where the entire burial chain appears to have designated places for each individual.

The Boz-Barmak burial ground is represented by six large kurgans (5, 8, 10, 11, 12 and 13), surrounded by smaller, less complex burials. One plausible interpretation is that the chain began with K10.1 and 10.2 as a couple, given both individuals buried were over 55 years old. Power may have then passed to K8 (female), followed by K5 (male), later to K13 (female) and finally to K12 and 11 (both females). This proposed sequence, together with the evidence of a single predominant Y chromosome lineage, could suggest that authority was transmitted through the paternal lineage, regardless of whether the successor was male or female.

We therefore hypothesize that the kinship relationships observed here might be concordant with a system of succession where power may have passed on not strictly from father to son, but to other relatives across the extended family, who were seen as the most influential at the time. Women in power could have included sisters, nieces, granddaughters, great-granddaughters, first cousins etc. of male individuals representing the local kinship. Such a speculative system of “Meritocratic Patrilineal Succession”, where meritocracy is sex or gender-neutral, finds parallels in ethnographic models of non-linear, or collateral, succession known in other pastoralist Inner Asian societies [32] and is potentially supported by the rich burials of other Scythian-Saka women [33, 34, 35], suggesting female authority was a feature of these cultures.

In conclusion, our analyses corroborate that Early Iron Age Saka people formed through extensive admixture between preceding populations from the Eurasian Steppe and various neighboring regions. The high diversity in mtDNA haplogroups and the predominance of a single Y chromosome lineage found in the Boz-Barmak individuals might reflect patrilocal practices. Findings from other Scythian groups and broader Eurasian steppe populations [2, 27] have been interpreted to show a similar pattern, leading to the hypothesis that the males may have remained within their natal groups, while females moved between communities. Taken together, our findings are in line with the suggestion that transmission of leadership among the Saka at Boz-Barmak may have had a complex nature, pointing to a sophisticated and adaptable sociopolitical organization.

## Methods

### DNA extraction and library preparation

DNA extractions and library constructions were performed in laboratories designed and dedicated only to ancient DNA (aDNA) research at the University of Vienna, wearing protective clothing and following aDNA laboratory best practices [36]. Specimens were treated with a sandblaster and ground to bone powder using a MixerMill (Retsch). DNA extraction was performed using an established protocol for aDNA [37], and Single-stranded DNA libraries were prepared [38]. We assessed quantity and quality using an Invitrogen QubitTM 4 Fluorometer and an Agilent 4150 TapeStation. Single-end sequencing was performed on the Illumina NovaSeq 6000 (SP SR100 XP workflow) at the Vienna BioCenter. Mitochondrial capture was performed using myBaits using the manufacturer’s instructions.

### Ancient DNA data processing and quality control

Raw FASTQ data from mitochondrial capture was trimmed with Cutadapt (version 5.1) [39] and raw reads from shotgun sequencing were trimmed using leehom (version v1.2.18) [40]. All the reads were then mapped to the hg19 human reference genome (GRCh37) http://ftp.1000genomes.ebi.ac.uk/vol1/ftp/technical/reference/ with the Burrows-Wheeler Aligner (BWA) aln algorithm (version 0.7.19-r1273) [41] using ancient DNA parameters. Polymerase chain reaction (PCR) duplicates were removed using Picard MarkDuplicates (version 3.1.1) [42]. Ancient DNA damage patterns were estimated using MapDamage2 (version 2.2.2) [43] and genotype calling was performed with bcftools mpileup (version 1.22) [44]. Modern human contamination in shotgun data was estimated with Calico (version 0.2) [45], and endogenous DNA content was calculated as proportion of uniquely mapped reads (MQ*>* 30) of total sequencing reads. Genotype imputation of autosomes and chromosome X was performed using the 1000 Genomes Project reference haplotype panel [46] via GLIMPSE1 (version v2.0.0) [47]. All scripts with parameters used for this study are available on the corresponding GitHub repository: https://github.com/Rymbekova/BozBarmak

### Genetic affinity and kinship analysis

Relatedness between individuals was determined with ancIBD (version 0.7) [21]. ancIBD works with the output from GLIMPSE1 - phased and imputed vcf files, which are then transformed to HDF5 files. Identical by descent (IBD) segments are called on HDF5 files for each autosome and combined. X chromosome IBD calling was done using ancIBDX command.

Principal Component Analysis for genetic affinities with broad Eurasian ancestry was done using PCAngsd (version v1.35) [48]. As the reference panel, vcf files of modern populations from Central Asia and Siberia, East Asia, South Asia and Europe from the Simons Genome Diversity Project (SGDP) [49] were merged together using bcftools merge. Genotype calling for two Kyrgyz individuals with the most coverage and one ancient Botai individual was performed with bcftools mpileup with -R option to restrict to the variants present in the reference panel and -f option for the hg19 reference fasta file. Ancient and modern Eurasians were then merged into one file using bcftools merge.

Principal Component Analysis for genetic affinities within the region during BA and IA was performed using the smartsnp R package (version 1.2.0) [50] As the reference panel, EIGENSTRAT files of BA and IA populations from Central Eurasia from the Allen Ancient DNA Resource (AADR) [51] merged with EIGENSTRAT files of Boz-Barmak individuals using EIGENSOFT package’s (version 8.0.0) convertf and mergeit commands [52]. PCA was performed using the reference populations and the Kyrgyz individuals were projected onto the resulting axes.

f3 statistic analysis was done using qp3Pop program (version 701) and qpAdm analysis was done using qpfstats (version 1002) and qpAdm (version 2050) programs from the AdmixTools package (version 8.0.2) [53]. The merged VCF file including Boz-Barmak individuals, ancient and modern-day Eurasians and Mbuti from the SGDP reference panel was converted to a eigenstrat format (.geno, .snp and .ind files) using a bash script convertVCFtoEigentstrat.sh https://github.com/joanam/scripts/tree/master.

### Runs of homozygosity

Using a bash script convertVCFtoEigentstrat.sh, the merged VCF file containing Boz-Barmak individuals was converted to the EIGENSTRAT format. Reference haplotype and metadata files for 1000 Genomes reference panel are provided together with the package, as well as Jupyter [54] notebooks for calling and plotting ROH using hapROH (version 0.64) [22].

### Haplogroup analysis

Mitochodrial DNA consensus calling was done with bcftools mpileup (version 1.22). Haplogroup assignment was conducted with haploGrouper (version 1.1) [55]. The program takes VCF files of mitochondrial DNA and Y chromosome as input. Pre-computed tree, locus and scores files for haplogroup assignment for both mitochondrial DNA and Y chromosome are provided with the program. Plots were created using custom R scripts (version 4.3.0) [56].

### Metagenomic assessment

For the metagenomic screening, raw FASTQ files were trimmed using trimmomatic (version 0.39) [57] and deduplicated with bbmap (version 39.06) clumpify [58]. Taxonomic classification was performed with Kraken2 using a database comprising all sequences from the NCBI nt database. The number of hits corresponding to relevant human-associated pathogens was assessed; however, due to the low number of hits, pathogen screening was not pursued further.

## Data availability

Sequence data are available in the European Nucleotide Archive (ENA) under accession number PRJEB97627.

## Acknowledgments

The computational work of this study was performed using the Life Science Compute Cluster (LiSC) of the University of Vienna.

## Author contributions

A.A., M. Hrivnyak, J.T.E. and K.K. provided sample material and archaeological background information. A. L.-L. and K.B. performed the ancient DNA sequencing under the supervision of O.C. A.R. performed data analysis and wrote the manuscript with input from all coauthors. M.Hämmerle performed the metagenomic screening. M.K. and P.G. supervised data analysis and writing the manuscript. A.A., J.T.E., M.Hrivnyak and R.P. provided help in writing the manuscript.

## Research funding

This project has been funded by the Vienna Science and Technology Fund (WWTF) [https://doi.org/10.47379/VRG20001] to M.K. The material is based upon work partially supported by the U.S. National Science Foundation under Grant No.2141844 awarded to J. T. E. and M. Hrivnyak. Any opinions, findings, and conclusions or recommendations expressed in this material are those of the author(s) and do not necessarily reflect the views of NSF. P. G is supported by European Research Council (ERC) under the SHADOWS project (grant agreement No. 101163057) and the Austrian Science Fund (FWF) [https://doi.org/10.55776/P36433]. Archaeological excavations of the site were funded by Bridge construction team of SE NK “Kyrgyz Temir Jolu”.

## Supplementary Information

### Supplementary Figures and Tables

**Table S1.** Results from mitochondrial DNA capture (Modern cont. = modern contamination).

| ID | Total reads | Aligned reads | Unique reads | Reads with quality > 30 | Modern cont.% |
| --- | --- | --- | --- | --- | --- |
| BB.K.1 | 11,367,543 | 4,583,220 | 33,044 | 32,761 | 2,1 |
| BB.Obj.2 | 12,724,850 | 9,666,832 | 32,932 | 32,220 | 9,9 |
| BB.K.4 | 11,349,329 | 127,446 | 3,350 | 3,123 | 0 |
| BB.K.6 | 11,586,891 | 9,594,249 | 33,072 | 32,553 | 2,7 |
| BB.K.8 | 10,961,268 | 468,843 | 4,181 | 3,922 | 5,1 |
| BB.K.9 | 13,770,508 | 11,805,917 | 33,069 | 32,293 | 1,7 |
| BB.K.10.1 | 11,325,963 | 8,327,901 | 32,906 | 32,175 | 3,3 |
| BB.K.10.2 | 11,286,571 | 4,902,714 | 17,088 | 16,077 | 0 |
| BB.K.11 | 11,100,431 | 7,029,265 | 33,071 | 32,777 | 1,3 |
| BB.K.12 | 13,870,541 | 9,847,270 | 33,068 | 32,007 | 3,1 |
| BB.K.14 | 12,297,707 | 8,741,174 | 33,070 | 32,620 | 2,3 |
| BB.K.15 | 11,676,636 | 7,814,114 | 33,071 | 32,440 | 2 |

**Table S2.** Total amount of runs of homozygosity (ROH) in centimorgans (cM) and the number of ROH fragments detected

| ID | Sum of ROH (cM) | Number of ROH fragments |
| --- | --- | --- |
| BB.K12 | 31.7 | 2 |
| BB.K15 | 9.5 | 1 |
| BB.K.6 | 6.4 | 1 |

**Figure S1.**
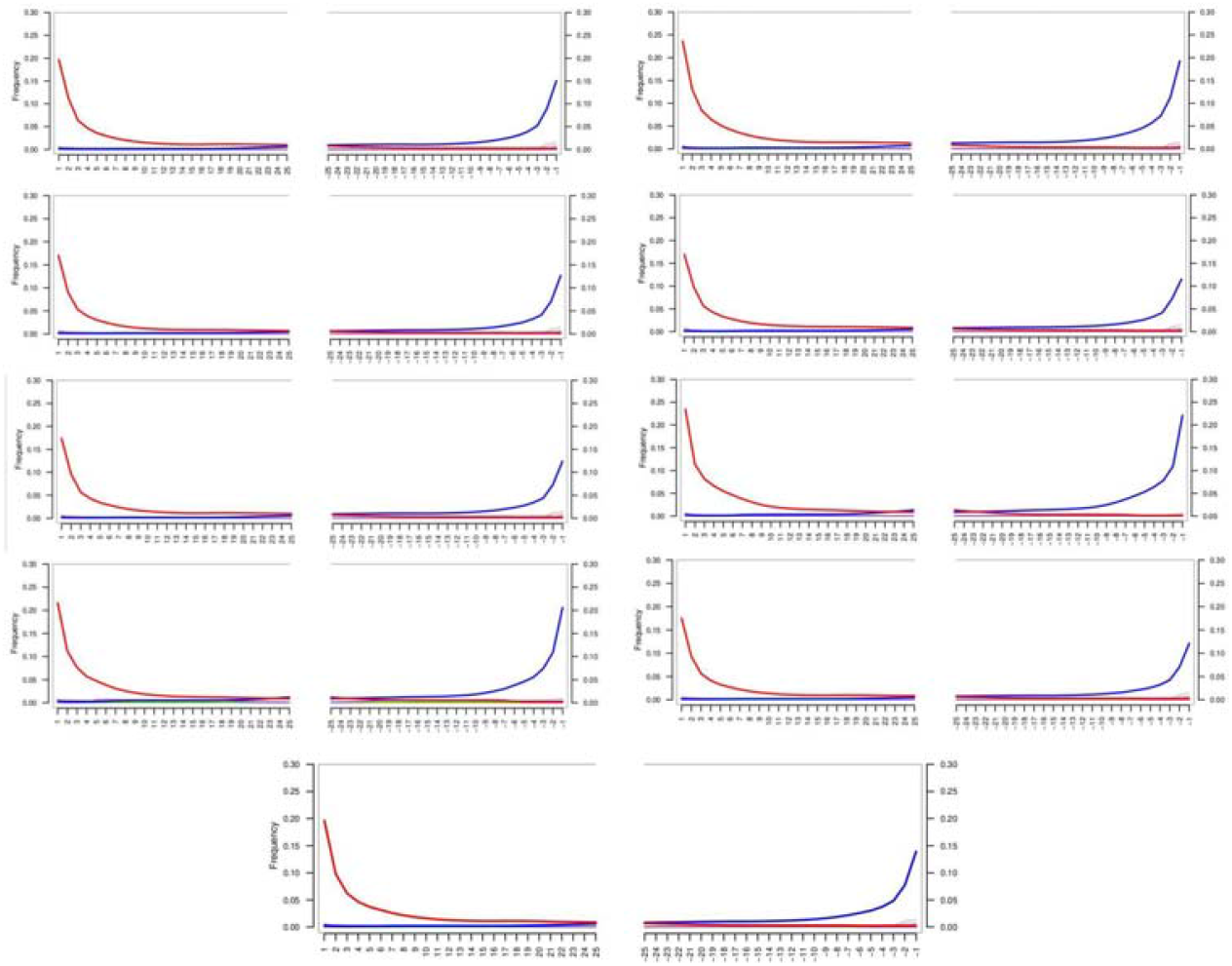
Ancient DNA damage patterns of nine Boz-Barmak individuals.

**Figure S2.**
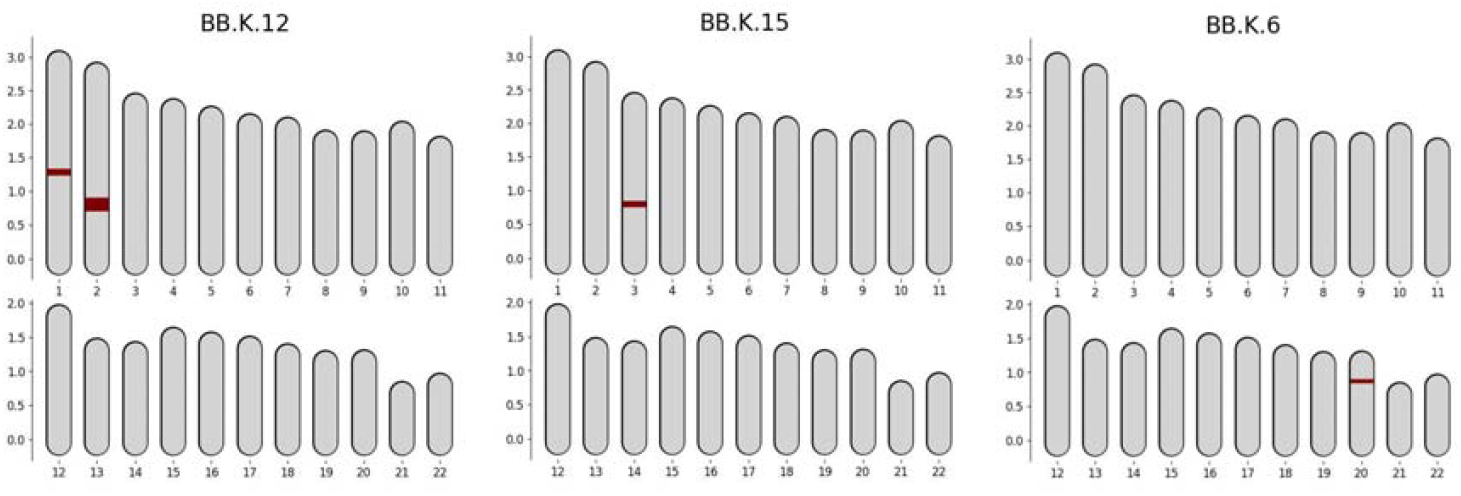
Runs of homozygosity detected in three Boz-Barmak individuals.

**Figure S3.**
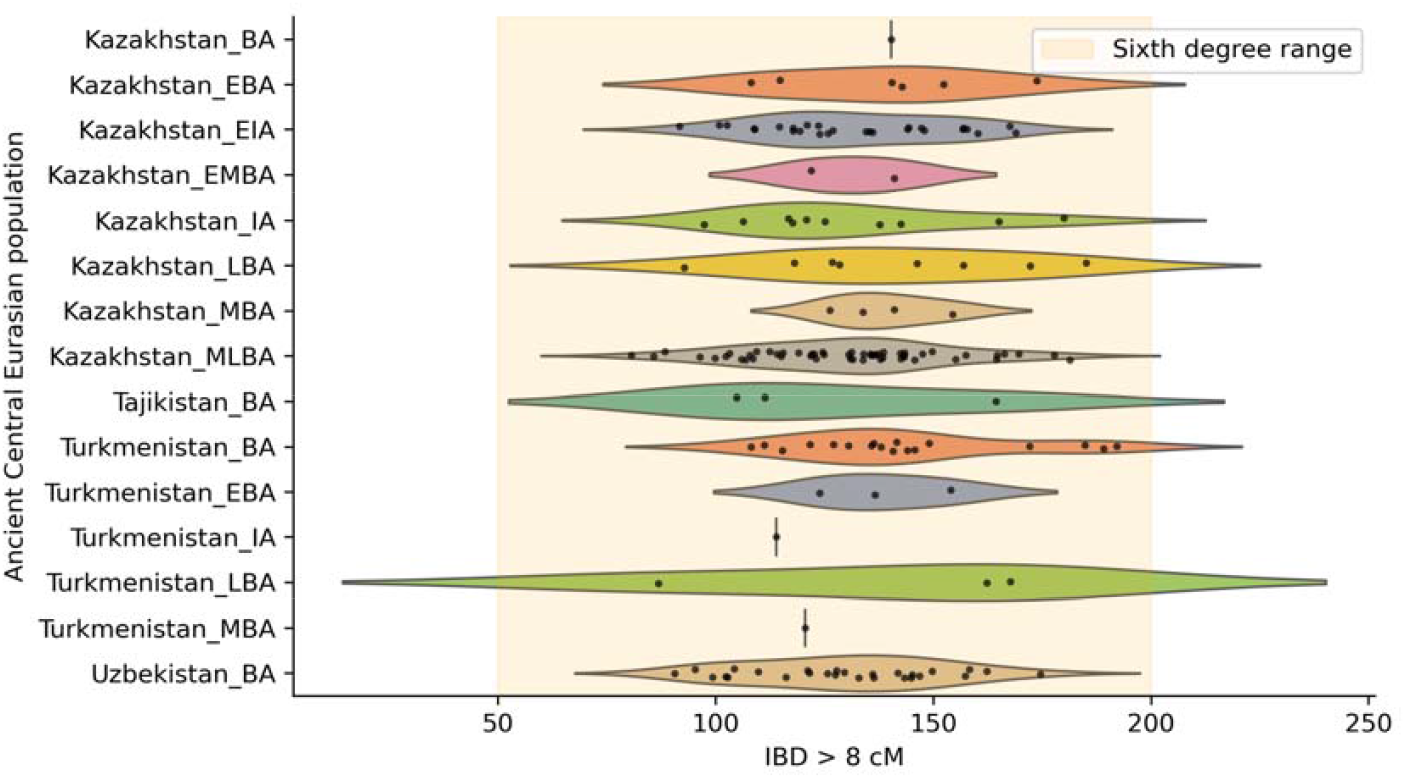
Identity by Descent (IBD) sharing in centimorgans (cM) between Boz-Barmak individuals and previously published populations, including fragments longer than 8 cM. Each point represents an individual of an ancient Central Eurasian population and its shared IBD with Boz Barmak individuals. (BA = Bronze Age, EBA = Early Bronze Age, MBA = Middle Bronze Age, MLBA = Middle Late Bronze Age, LBA = Late Bronze Age, IA = Iron Age, EIA = Early Iron Age)

## References

1. Haak, W. et al. Massive migration from the steppe was a source for Indo-European languages in Europe. Nature 522, 207–211. ISSN: 0028-0836. http://www.nature.com/doifinder/10.1038/nature14317 (2015).

2. Allentoft, M. E. et al. Population genomics of Bronze Age Eurasia. Nature 522, 167. ISSN: 0028-0836. http://www.nature.com/doifinder/10.1038/nature14507 http://dx.doi.org/10.1038/nature14507 http://10.0.4.14/nature14507 https://www.nature.com/ articles/nature14507#supplementary-information (June 2015).

3. Gnecchi-Ruscone, G. A. et al. Ancient genomic time transect from the Central Asian Steppe unravels the history of the Scythians. Science Advances 7, eabe4414. 10.1126/sciadv.abe4414 (July 2025).

4. Narasimhan, V. M. et al. The formation of human populations in South and Central Asia. Science 365, eaat7487. 10.1126/science.aat7487 (Sept. 2019).

5. Jeong, C. et al. The genetic history of admixture across inner Eurasia. Nature Ecology & Evolution 3, 966–976. ISSN: 2397-334X. 10.1038/s41559-019-0878-2 (2019).

6. Akishev, K.A. & Kushaev, G.A. Ancient history of Saka and Usun of the Ili river valley. Publishing House of the Academy of Sciences of the Kazakh SSR. (in Russian) (1963).

7. Davis-Kimball, J., Bashilov, V. & Yablonsky, L. Nomads of the Eurasian Steppes in the Early Iron Age. (Zinat Pr, 1995).

8. Khasanov, A. A social history of the Scythians. Moscow: Nauka (in Russian) (1975).

9. Gulyayev V.I. Scythians: The rise and fall of a great kingdom. (in Russian) (2006).

10. Kumar, V., Wang, W., Zhang, J. & et. al. Bronze and Iron Age population movements underlie Xinjiang population history. Science 376, 62–69. eprint: https://www.science.org/doi/pdf/10.1126/science.abk1534. https://www.science.org/doi/abs/10.1126/science.abk1534 (2022).

11. Jeong, C. et al. A Dynamic 6,000-Year Genetic History of Eurasia’s Eastern Steppe. Cell 183, 890–904. ISSN: 0092-8674. https://www.sciencedirect.com/science/article/pii/S0092867420313210 (2020).

12. Unterlander, M., Palstra, F., Lazaridis, I. & Pilipenko, A. Ancestry and demography and descendants of Iron Age nomads of the Eurasian Steppe. Nat Commun. 8, 14615 (2017).

13. Sadykov, T., Caspari, G. & Blochin, J. Kurgan Tunnug 1 — New Data on the Earliest Horizon of Scythian Material Culture, Journal of Field Archaeology, DOI:10.1080/00934690.2020.1821152 (2020).

14. Damgaard, P., Marchi, N., Rasmussen, S. & et al. 137 ancient human genomes from across the Eurasian steppes. Nature 557, 369–374 (2018).

15. Jarve, M. et al. Shifts in the Genetic Landscape of the Western Eurasian Steppe Associated with the Beginning and End of the Scythian Dominance. Curr Biol. 29, 2430–2441 (2019).

16. Koryakova, L. & Epimakhov, A. The Urals and Western Siberia in the Bronze and Iron Ages. 10.1017/cbo9780511618451 (2007).

17. Zaitseva, G., Chugunov, K., Alekseev, A., Dergachev, V. & et. al. Chronology of Key Barrows Belonging to Different Stages of the Scythian Period in Tuva (Arzhan-1 and Arzhan-2 Barrows) Radiocarbon. 49, 645–658 (2007).

18. Kubarev, V.D., Shulga, P.I. Pazyryk culture (Chuya and Ursul kurgans). Barnaul, 282 p. ISBN 978-5-7904-0684-3 (in Russian) (2007).

19. Rakhimov, N.T. Family and Marital Relations of the Ancient Saka (Toward the History of Family Law of the Tajiks). Bulletin of the Tajik State University of Law, Business and Politics. Series of Social Sciences, 4, 93–100 (in Russian) (2016).

20. Piyankov, I.V. History of Persia by Ktesiy and Central Asiatical Satrapies Referring to Ahemenids at the End of the 5^th^ century BCE. Bulletin of Ancient History, 2, 84–91 (in Russian) (1965).

21. Ringbauer, H., Huang, Y., Akbari, A. & et al. Accurate detection of identity-by-descent segments in human ancient DNA. Nat Genet. 56, 143–151 (2024).

22. Ringbauer, H., Novembre, J. & Steinrucken, M. Parental relatedness through time revealed by runs of homozygosity in ancient DNA. Nat Commun. 12 (2021).

23. Harney, E. et al. Assessing the performance qpAdm: a statistical tool for studying population admixture. Genetics 4, iyaa045. 10.1093/genetics/iyaa045 (2021).

24. Lerner, A. Iron Age Nomads of the Urals: Interpreting Sauro-Sarmatian and Sargat Identities. UMI ProQuest, Ann Arbor (2006).

25. Koryakova, L. & Daire, M. Burials and Settlements at the Eurasian Crossroads: Joint Franco-Russian Project. (2000).

26. Matveeva, N.P. Socio-Economic Structure of the Population of Western Siberia in the Early Iron Age. Novosibirsk: Russian Academy of Sciences-Siberian Division and Nauka (in Russian) (2000).

27. Krzewinska, M. et al. Ancient genomes suggest the eastern Pontic-Caspian steppe as the source of western Iron Age nomads. Science Advances 4, eaat4457, 10.1126/sciadv.aat4457 (2018).

28. Der Sarkissian, C. et al. Genographic Consortium, Ancient DNA reveals prehistoric gene-flow from Siberia in the complex human population history of North East Europe. PLOS Genet. 9, e1003296 (2013).

29. Juras, A.. et al. Diverse origin of mitochondrial lineages in Iron Age Black Sea Scythians. Sci. Rep. 7, 43950 (2017).

30. Kyzlasov, I.L. Prot-Turkic dwellings. Survey of Sayan-Altai antiquities. Moscow-Samara, 96 p.: ill. (in Russian) (2005).

31. Kubarev, V.D. Kurgans of Saylyugem. Novosibirsk, 224 p. ISBN 5-02-030225-2 (in Russian) (1992)

32. Barfield, T. J. The Perilous Frontier. Nomadic Empires and China. Cambridge (1989)

33. Polosmak, N. The Horsemen of Ukok. Novosibirsk: INFOLIO-press, p.337, ISBN 9785041538415 (in Russian) (2000).

34. Simpson, S. J., & Pankova, S. Scythians: warriors of ancient Siberia (2017).

35. Gulyaev, V., Volodin, A., Shevchenko, A. Excavations of Kurgan number 9 at the Devitsa V Burila Ground. Archaeological Research in the Central Chernozem Region 2019. Lipetsk, Voronezh, p. 60–64 (in Russian) (2020).

36. Childebayeva A, Zavala EI. Review: Computational analysis of human skeletal remains in ancient DNA and forensic genetics. iScience. 2023 Oct 4;26(11):108066. doi: 10.1016/j.isci.2023.108066. PMID: 37927550; PMCID: PMC10622734.

37. Dabney, J. et al. Complete mitochondrial genome sequence of a Middle Pleistocene cave bear reconstructed from ultrashort DNA fragments. Proceedings of the National Academy of Sciences 110, 15758–15763. https://www.pnas.org/doi/abs/10.1073/pnas.1314445110 (2013).

38. Kapp, J., Green, R. & Shapiro, B. A Fast and Efficient Single-stranded Genomic Library Preparation Method Optimized for Ancient DNA. J Hered. 112, 241–249 (2021).

39. Martin, M. Cutadapt removes adapter sequences from high-throughput sequencing reads. EMBnet.journal 17, 10–12. ISSN: 2226-6089 (2011).

40. Renaud, G., Stenzel, U. & Kelso, J. leeHom: adaptor trimming and merging for Illumina sequencing reads. Nucleic Acids Research 42, e141–e141. ISSN: 0305-1048. e-print: https://academic.oup.com/nar/article-pdf/42/18/e141/17423287/gku699.pdf. 10.1093/nar/gku699 (Aug. 2014).

41. Li, H. & Durbin, R. Fast and accurate short read alignment with Burrows-Wheeler transform. Bioinformatics. 25, 1754–1760 (2009).

42. Broad Institute. Picard Tools Website. Accessed: 2025-06-25. 2025. http://broadinstitute.github.io/picard/.

43. Ginolhac, A., Rasmussen, M., Gilbert, M. T. P., Willerslev, E. & Orlando, L. mapDamage: testing for damage patterns in ancient DNA sequences. Bioinformatics 27, 2153–2155. ISSN: 1367-4803 (June 2011).

44. Danecek, P. et al. Twelve years of SAMtools and BCFtools. Gigascience. 10 (2021).

45. Skoglund, P. Estimating DNA contamination GitHub repository. Accessed: 2025-06-24. 2025.https://github.com/pontussk/calico.

46. 1000 Genomes Project Consortium et al. A global reference for human genetic variation. Nature. 526, 68–74 (2015).

47. Rubinacci, S., Hofmeister, R., Sousa da Mota, B. & Delaneau, O. Imputation of low-coverage sequencing data from 150,119 UK Biobank genomes. Nat Genet. 55, 1088–1090 (2023).

48. Meisner, J. & Albrechtsen, A. Inferring Population Structure and Admixture Proportions in Low-Depth NGS Data. Genetics 210, 719–731. ISSN: 1943-2631. eprint: https://academic.oup.com/ genetics/article-pdf/210/2/719/49481227/genetics0719.pdf. 10.1534/genetics.118.301336 (Aug. 2018).

49. Mallick, S. et al. The Simons Genome Diversity Project: 300 genomes from 142 diverse populations. Nature. 538, 201–206 (2016).

50. Herrando-Pérez, S., Tobler, R., & Huber, C. D. smartsnp, an r package for fast multivariate analyses of big genomic data. Methods in Ecology and Evolution, 12, 2084–2093. 10.1111/2041-210X.13684 (2021).

51. Mallick, S. and Reich, D. The Allen Ancient DNA Resource (AADR): A curated compendium of ancient human genomes. 10.7910/DVN/FFIDCW (2023).

52. Patterson, N., Price, A.L., Reich, D. Population Structure and Eigenanalysis. 10.1371/journal.pgen.0020190 (2006).

53. Patterson, N. et al. Ancient admixture in human history. Genetics. 192, 1065–93 (2012).

54. Kluyver, T., Ragan-Kelley, B., Perez, F. & et.al. Jupyter Notebooks – a publishing format for reproducible computational workflows in Positioning and Power in Academic Publishing: Players, Agents and Agendas (eds Loizides, F. & Schmidt, B.) (2016), 87–90.

55. Jagadeesan, A. et al. HaploGrouper: a generalized approach to haplogroup classification. Bioinformatics 37, 570–572. ISSN: 1367-4803. eprint: https://academic.oup.com/bioinformatics/article-pdf/37/4/570/50359810/btaa729.pdf. 10.1093/bioinformatics/btaa729 (Aug. 2020).

56. R Core Team. R: A Language and Environment for Statistical Computing R Foundation for Statistical Computing (Vienna, Austria, 2021). https://www.R-project.org/.

57. Bolger, A. M., Lohse, M. & Usadel, B. Trimmomatic: A flexible trimmer for Illumina sequence data. Bioinformatics 30, 2114–2120 (2014).

58. Bushnell, B. BBMap: A Fast, Accurate, Splice-Aware Aligner.

